# The role of iron uptake systems in the pathogenesis of colistin-resistant hypervirulent *K.pneumoniae* infections

**DOI:** 10.1101/677492

**Authors:** Ozlem Dogan, Cansel Vatansever, Nazli Atac, Ozgur Albayrak, Sercin Karahuseyinoglu, Ozgun Ekin Sahin, Bilge Kaan Kilicoglu, Atalay Demiray, Onder Ergonul, Mehmet Gönen, Fusun Can

## Abstract

Here we proposed the hypothesis that hypervirulent colistin resistant *K.pneumoniae* (ColR-Kp) exhibit high number of virulence factors and have enhanced survival capacity against neutrophil activity.

We studied virulence genes of ColR-Kp isolates and neutrophil response in 142 patients with invasive infections.

The patients infected with hypervirulent ST101 and ST395 ColR-Kp had higher 30-day mortality (58%, p=0.005 and 75%, p=0.003, respectively. The yersiniabactin biosynthesis gene (ybtS) and ferric uptake operon associated gene (kfu) were significantly higher in ST101 (99%, p=<0.001) and in ST395 (94%, p<0.012). Being in ICU (OR: 7.9; CI: 1.43-55.98; p=0.024), kfu (OR:27.0; CI:5.67-179.65; p<0.001) and ST101 (OR: 17.2; CI: 2.45-350.40; p=0.01) were found to be predictors of 30-day mortality. The uptake of kfu^+^-ybtS^+^ ColR-Kp by neutrophils was significantly higher than kfu^-^-ybtS^-^ ColR-Kp (78% vs 65%, p<0.001). However, kfu^+^-ybtS^+^ ColR-Kp were more resistant to the killing activity of neutrophils than negative ones (7.90 vs 4.22; p=0.001). The kfu^+^-ybtS^+^ ColR-Kp stimulated excessive NET formation while the NET’s against kfu^-^-ybtS^-^ ColR-Kp were weak and rare.

Iron uptake systems enhance successful survival of *K.pneumoniae* against neutrophil phagocytic defense, and stimulate excessive NET formation. The drugs targeted to iron uptake systems would be a promising approach for treatment of hypervirulent *K.pneumoniae* infections.

## Introduction

Colistin resistant hypervirulent *Klebsiella pneumonia* (HvKp) infections are one of the emerging threats in public health because of high fatality rates (1–3). The ST101 and ST395 clones of *K.pneumonia* are known as hypervirulent clones (4–7), and reported to be significant predictors of the mortality (8). The leading virulence factors of HvKp are mostly associated with capsular serotype, muco-viscosity, iron uptake systems and allantoin metabolism (9–11). Enhanced adhesion and attachment by fimbria and non-fimbrial-structures promote pathogenicity of *K.pneumoniae* as well (9, 12).

Iron uptake system is essential for survival and dissemination of pathogens during infections. These systems have also a significant effect on host inflammatory response (13). The neutrophils as the important cells of the immune defense, kill pathogens by engulfment or release of extracellular traps (NETs) (14). The function of NETs is to trap bacteria and promote extracellular killing by minimizing damage to host cells (14). A previous study reported that low phagocytic activity of neutrophils contributes to the success of carbapenem-resistant HvKp ST258 clone (15). However, our knowledge on immune escape mechanisms of colistin-resistant hypervirulent *K.pneumoniae* (ColR-HvKp) is very limited (10, 11). By this study, we aimed to describe the role of the major virulence factors of ColR-HvKp and their interaction with the neutrophils. Our results will provide an insight to depict the pathogenesis of the HvKp infection.

## Methods

### Study population and Data Collection

The patients diagnosed with colistin resistant *K.pneumoniae* infection between January 2015 and May 2018 were included in the study. A study protocol reviewing patient’s demographic data, underlying diseases, type of infection, isolation site, blood biochemical parameters, predisposing factors such as having operation within last one month, intensive care unit admission (ICU), type of antimicrobial agents used for empirical and agent-specific therapy, duration of colistin therapy before isolation of colistin-resistant isolates, and carbapenem resistance was used. The patients were followed up for fatality for 30 days after hospital admission. Exclusion criteria were missing key data, subsequent episodes of the same patient.

### Microbiological and Molecular Studies

Colistin resistance was studied by broth microdilution and breakpoint for resistance was set to >2 mg/L (16). Carbapenemase genes of OXA-48, NDM-1, KPC were examined by multiplex-PCR, and amplicons were sequenced (17). The *mcr-1* was screened by PCR described by Liu et al (18).

Genotyping of the isolates was carried out by MLST comparing seven housekeeping genes (*phoE, gapA, rpoB, tonB, inf, mdh, and pgi*) according to the protocol published on the Institute of Pasteur website (http://bigsdb.pasteur.fr/klebsiella/klebsiella.html). ST types were determined using Applied Maths Bionumerics V7.6 software.

Virulence genes of type-1 and type-3 adhesins (*FimH-1, mrkD*), enterobactin biosynthesis (*entB*), aerobactin receptor (*iutA*), yersiniabactin receptor (*fyuA*), yersiniabactin biosynthesis (*ybtS*), ferric uptake operon associated gene (*kfu*), regulator of mucoid phenotype A (*rmpA*), capsule type 1 (*magA*), capsule type2 (*K2Wzy*), capsule type 5 (*K5wzx*), outer core lipopolysaccharide biosynthesis (*wabG*) and allantoin metabolism (*allS*) were screened by PCR using primers described previously (19)

### Phagocytosis assays

For phagocytosis assays, 10 *ybtS*^+^-*kfu*^+^, eight *ybtS*^−^-*kfu*^-^, two *ybtS*^−^*kfu*^+^, and one *ybtS*^+^*kfu*^-^ isolates were selected. K. *pneumoniae* ATCC 700831 and S. *epidermidis* ATCC 35984 were used as controls. Human neutrophils were separated from peripheral blood by density gradient centrifugation using Histopaque (Sigma–Aldrich, Germany) according to the manufacturer’s instructions. Neutrophil purity was determined by Flow Cytometry (BD Biosciences, USA) using mouse anti-human CD15-PE (Bechman-Coulter, USA) antibody. K. *pneumoniae* isolates were stained with BacLight 488 (Thermo Scientific, USA) with slight changes to manufacturer’s instructions. For phagocytosis, 2 x 10^7^ neutrophils were incubated with bacterial suspension containing 3 x 10^8^ bacteria for 30 minutes at 37 ^0^C. Phagocytosis was stopped by adding 1ml of ice-cold PBS into tubes. A portion of each sample was stained with Mouse-Anti Human CD15-PE (Beckman Coulter, USA) and run under BD Accuri C6 Flow Cytometer. The internalized and/or surface attached bacteria were determined as CD15^+^BacLight 488+ cells whereas free bacteria were determined as only BacLight 488^+^ Cells. Phagocytic Index (Ph Index) was calculated by [(Initial bacterial count X Ph%)/100]. For viability testing, neutrophils were lysed with dH_2_O for 20 minutes and cultured on tryptic soy agar by 10 fold dilutions. After overnight incubation, colonies were counted and the survival index was calculated by [(Colony count per ml/Ph Index) X100].

### Detection of Neutrophil Extracellular Traps

Two *ybtS*^+^-*kfu*^+^ and two *ybtS*^-^-*kfu*^-^ isolates were selected for NETosis experiments. Neutrophils (2 x 10^5^ cells) incubated for 1 hour at 37^0^C for attachment to the surface. After incubation, 6 x 10^6^ bacteria were added on neutrophils and incubated 90 minutes at 37^0^C for NET generation (1:30). A portion of each cell was fixed and permeabilized with 4% BSA and 0,2 % Triton X-100. After blocking, the cells were stained with Mouse Anti-Human Myeloperoxidase (Santacruz, Germany) and Rabbit Anti-Human Histone-H3 (Abcam, USA) antibodies for one hour. Rabbit Anti-mouse Alexa-Fluor 594 (Biolegend, USA) and Goat Anti-rabbit Alexa-Fluor 488 (Thermo Scientific, USA) were used as secondary antibodies. Fluoreshield medium with DAPI (Abcam, USA) was used for mounting and analyses were performed under confocal microscope (Leica dmi8/Sp8, Germany). K. *pneumoniae* ATCC700831 was used as control. The remaining part was assessed for the viability of the bacteria after NET formation. Cell suspensions were cultured on tryptic soy agar by 10 fold dilutions and colony count/ml was recorded.

### Statistical analysis

Statistical analysis was performed using the statistical software package R. In univariate analyses, we used Wilcoxon rank-sum test for continuous covariates and Fisher’s exact test for discrete covariates. In multivariate analyses, logistic regression was performed using the variables that were detected to be significant in univariate analyses. All the results of statistical analysis are available at the supplementary file (https://midaslab.shinyapps.io/klebsiella_pneumoniae_virulence_analysis/)

## Results

In this study, 142 patients with colistin resistant *K.pneumoniae* infection out of 710 (20%) were analyzed (Figure 1). In study group, 84% of the patients stayed in ICU, bacteremia was detected among 43% of the patients and 47% of them had ventricular associated pneumonia (VAP). The median age of the patients was 61 and 58% of the patients were male. The 30-day mortality was 51%.

**Figure 1.**
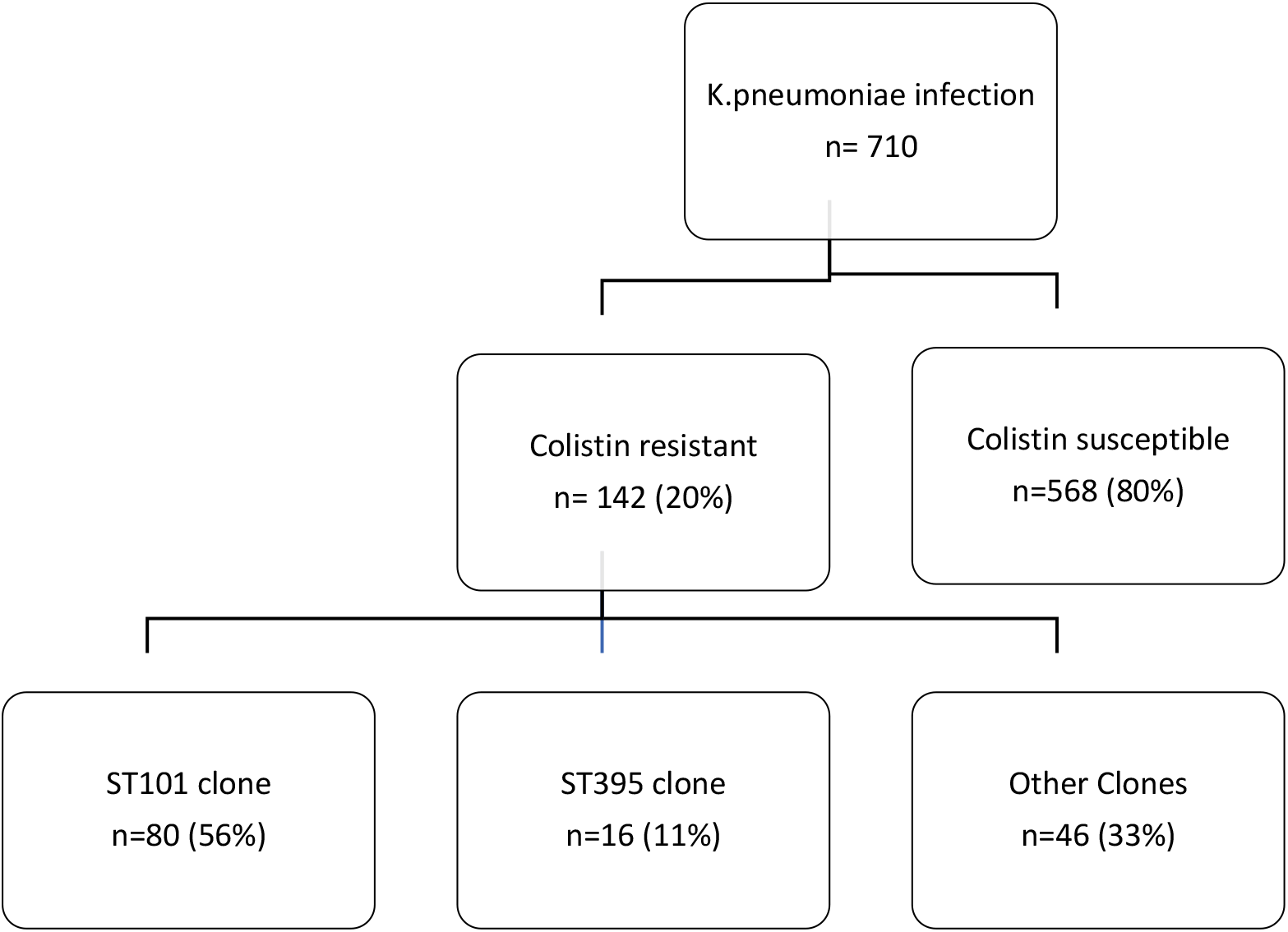
Distribution of the hypervirulent clones in *Klebsiella pneumoniae*.

All the isolates were resistant to colistin, with MICs between 4 and 256mg/L. The majority of ColR-Kp belonged to ST101 (56%) and ST395(11%) HvKp, and the others distributed to various ST clones (minimum spanning tree in supplement) The patients infected with ST101 ColR-Kp had more VAP (56%, p=0.006), had higher 30-day mortality rate (58%p=0.005) than other clones. The mortality rate among ST395 type *K.pneumoniae* infected patients was 75% (p=0.003) (Table 1).

**Table 1:** Clinical characteristics of the patients infected with colistin resistant *K.pneumoniae*

| Patient | Total (n=142)<br>Median (range) | ST101 n=80<br>n (%) | ST395 n=16<br>n (%) | Others*(n=46)<br>n (%) |
| --- | --- | --- | --- | --- |
| Age | 61 (0-91) | 63 (0-86) | 62 (30-84) | 53 (0-91) |
| Female Gender | 60 (42) | 34 (43) | 7 (44) | 19 (41) |
| Bacteremia | 61 (43) | 33 (41)<br>P=0.852 | 10(63)<br>P=0.147 | 18 (39) |
| VAP | 67 (47) | 44 (55)<br>P=0.009 | 9(56)<br>P=0.079 | 14 (30) |
| Mortality | 95 (67) | 61 (76)<br>P=0.002 | 12(75)<br>P=0.082 | 22 (48) |
| 30-day mortality | 72(51) | 46 (58)<br>P=0.005 | 12(75)<br>P=0.003 | 14 (30) |
| Being in ICU | 119 (84) | 68 (86)<br>P=0.323 | 15(94)<br>P=0.261 | 36 (78) |

\*Non-ST101 and Non-ST395 ColR-Kp

Among virulence factors, ferric uptake operon associated gene (*kfu*) and yersiniabactin (ybtS) components of iron uptake systems were found to be significantly higher in ST101 and ST395 ColR-Kp compared to the other clones. The *ybtS* and *kfu* positivity were 99% in ST101 (p=<0.001) and 94% in ST395 clones (p<0.012). The mucoid type associated gene (*rmpA*) and fimH type adhesin were also significantly higher in ST101 with the percentage of 89%, (p=0.005) and 99 %, (p=0.024), respectively.

The carriage of OXA-48 carbapenemase was significantly higher (95%, p=0.003) in ST101 than the other clones (76%), however it was found to be very low (31%, p=0.002) in ST395 clone. On the contrary, NDM-1 production was significantly higher in ST395 (88%, p<0.001), and lower in ST101 (4%, p<0.001) than the other clones(30%) (Table 2).

**Table 2:** The virulence factors and carbapenemase types in the ColR-Kp ST101 and ST395 clones

|  | Mucoïd type and Capsule type<br>n (/%) |  |  |  | Iron metabolism<br>n (/%) |  |  |  |  | Adhesins<br>n (/%) |  | LPS synt<br>n (/%) | Allantoin<br>met n (/%) | Carbapenemase<br>type n (/%) |  |
| --- | --- | --- | --- | --- | --- | --- | --- | --- | --- | --- | --- | --- | --- | --- | --- |
|  | *RmpA* | *MagA* | *K2Wzy* | *K5wzx* | *FyuA* | *Kfu* | *lutA* | *ybtS* | *entB* | *mrkD* | *FimH* | *WabG* | *AllS* | OXA-48 | NDM-1 |
| ST101<br>N=80 | 71<br>(89) | 6<br>(8) | 31<br>(39) | 0 | 79<br>(99) | 79<br>(99) | 4<br>(5) | 79<br>(99) | 80<br>(100) | 79 (99) | 79<br>(99) | 80 (100) | 0 | 76 (95) | 3 (4) |
| p | 0.005 | 1 | 0.707 |  | 0.553 | <0.001 | 0.285 | <0.001 |  | 0.059 | 0.024 | 0.365 |  | P=0.00 | P<0.001 |
| ST395<br>n=16 | 14<br>(88) | 0 | 9<br>(56) | 0 | 15<br>(94) | 15<br>(94) | 1<br>(6) | 15<br>(94) | 15<br>(100) | 15 (94) | 16<br>(100) | 16 (100) | 0 | 5<br>(31) | 14<br>(88) |
| p | 0.194 | 0.565 | 0.401 |  | 1 | 0.012 | 1 | 0.012 |  | 1 | 0.315 | 1 |  | P=0.002 | P<0.001 |
| Others<br>n=46* | 32<br>(68) | 23<br>(49) | 20<br>(43) | 0 | 45<br>(96) | 28<br>(60) | 5 (11) | 28<br>(60) | 47<br>(100) | 43 (92) | 42 (89) | 46<br>(98) | 0 | 36<br>(77) | 14<br>(30) |

\*Non ST101 and NonST395 ColR-Kp

In univariate analysis, being in ICU (OR: 4.3; CI: 1.42-16.04; p=0.005), presence of *ybtS* (OR: 3.0; CI: 1.01-10.02; p=0.034 and *kfu* (OR: 3.9; CI: 1.27-14.63; p=0.009) were found to be associated with 30-day mortality. In multivariate analysis, being in ICU (OR: 7.9; CI: 1.43-55.98; p=0.024), *kfu* (OR:27.0; CI:5.67-179.65; p<0.001) and ST101 (OR: 17.2; CI: 2.45-350.40; p=0.01) were found to be the predictors of 30-day fatality (supplement).

The phagocytosis experiments showed that *ybtS* and *kfu* positive ColR-Kp were internalized at higher rates (median=78%) while negative isolates exhibited low phagocytosis rates (median=65%) after 30 minutes of interaction with neutrophils (p<0.001, Figure 2). The phagocytosis rates of *S.epidermidis* and *K.pneumomniae* ATCC controls were 89% and 73%, respectively. Survival of *kfu*^+^-*ybtS*^+^ positive ColR-Kp was significantly higher than negative isolates with median survival index of 7.90 (range:3.29-13.13) vs 4.22 (range:0.36-5.64), respectively (p=0.001). The survival index of S.epidermidis and *K.pneumoniae* controls were 0.64 and 1.89, respectively (Figure2). The survival index of two *ybtS*^−^*kfu*^+^ isolates were 12.04 and 12.13, and it was 5.13 in one *ybtS*^+^*kfu*^-^ isolate.

**Figure 2.**
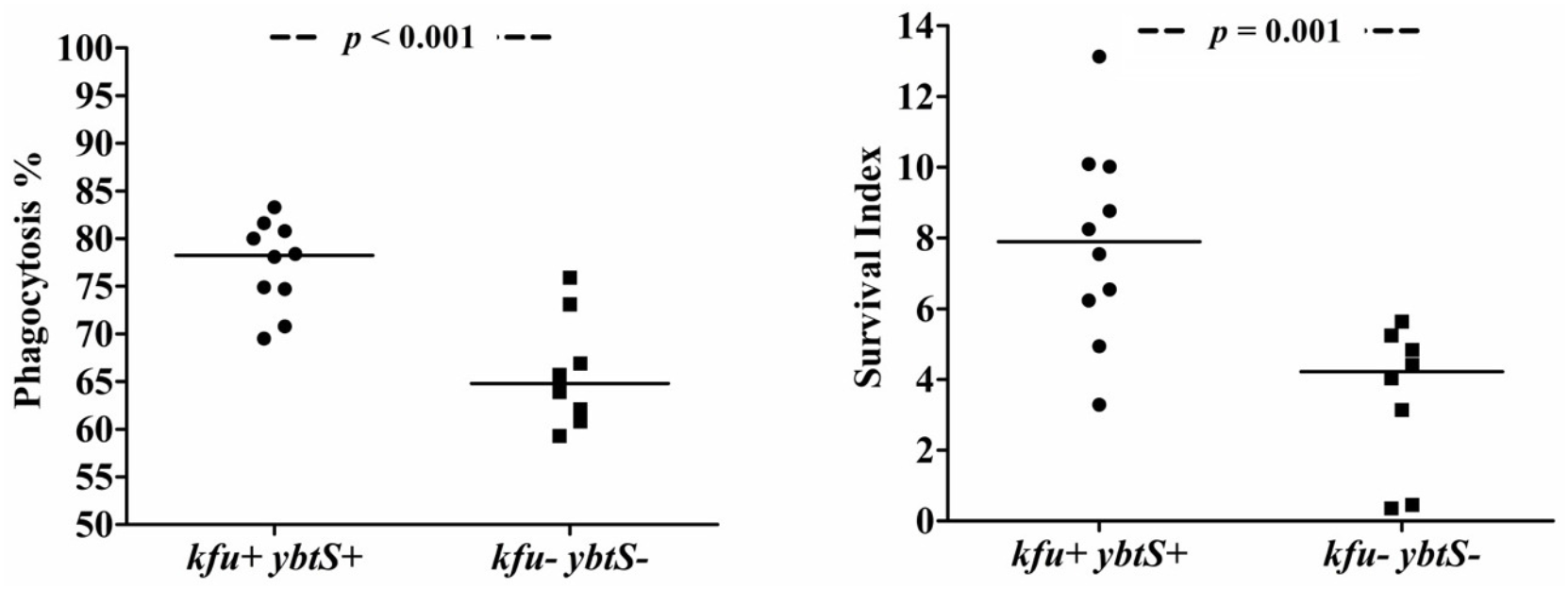
The phagocytosis of ColR-Kp by neutrophils. Phagocytosis rate of *kfu*^+^-*ybtS*^+^ and *kfu*^-^-*ybtS*^-^ isolates (A); Survival of *kfu*^+^-*ybtS*^+^ and *kfu*^-^-*ybtS*^-^ isolates after being phagocytosed by neutrophils (B)

Among 21 isolates, four was in ST101 clone. The median phagocytosis rate 80% was found to be and the survival index was 8.51.

After NETosis, the mean colony count of two *kfu*^+^-*ybtS*^+^ isolates was 5.50X10^6^ and it was 4.05×10^6^ for kfu^-^-*ybtS*^-^ strains. The colony count of ATCC K.pneumoniae was 4.3×10^6^. Confocal microscopy study showed that the *kfu*^+^-*ybtS*^+^ isolate stimulated abundant NET formation with excessive release of chromatin granular content to the extracellular area. However, the NET’s against *kfu*^-^-*ybtS*^-^ negative ColR-Kp were weak and seen only in few areas (Figure 3).

**Figure 3.**
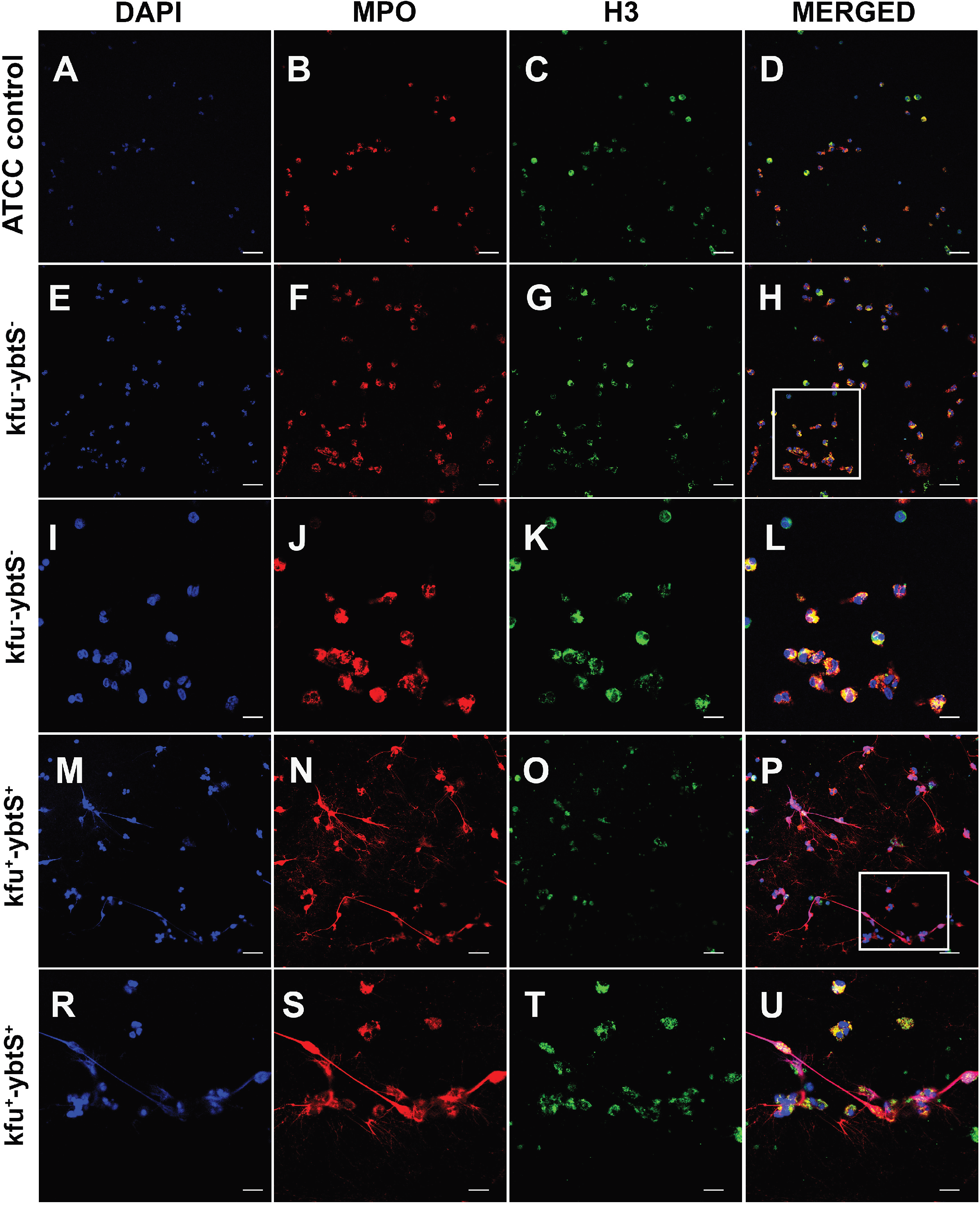
Confocal microscopic images of NETs. The samples were stained consecutively with myeloperoxidase (MPO, red) and histone 3 (H3, green). The nuclei were counterstained with DAPI (blue). Neutrophils were seen intact with K. pneumonia ATCC 700831 control(A-D). The *kfu*^-^-*ybtS*^-^ isolates depicted rare and weak NET formation (E-L). The rectangular area in image H was magnified in images I-L. The *kfu*^+^-*ybtS*^+^ isolates showed abundant NET formation with excessive histone and MPO release in extracellular matrix (M-U). The rectangular area in image P was magnified in images I-L. *Bars: A-H, M-P= 25 μm; I-L, R-U= 10 μm*.

## Discussion

ColR-HvKP infections are usually fatal because of limited therapeutic options due to extensive drug resistance and successful immune escape mechanisms of these pathogens. Alternative approaches are urgently needed in order to prevent and treat infections. One of the most promising strategies is inhibition of virulence factors. Here, we demonstrated the role of iron uptake systems in virulence of ColR-HvKP ST101 and ST395 clones.

We determined significantly higher iron uptake associated gene (*kfu*) and yersiniabactin (ybtS) positivity in ST101 (99%) and ST395 (94%) isolates (p<0.001, and p=0.012, respectively). These genes were found to be associated with 30-day mortality. Yersiniabactin type siderophore is encoded in high pathogenicity island which is responsible for high mortality and dissemination of infections (20, 21). It was also reported to be associated with pulmonary infections (3, 22). Holden et al. demonstrated that during pneumonia siderophores stabilizes HIF-1α and increases bacterial dissemination to the spleen (13). Lawlor et al. reported that the acquisition of yersiniabactin is an important step in the evolution of virulent *K.pneumoniae* (3). The kfu system was found to be associated with invasive infections and increased virulence in mice (23). In our study, multivariate analysis showed that *kfu* predicts 30-day mortality (OR: 27; CI: 5.67-179.56; p<0.001) and is a predictor of belonging ST101 clone (OD:20.3; CI: 2.17-484.56; p=0.018).

Another important disease strategy of hypervirulent clones is immune evasion from the innate response (24). The interaction of siderophores with host cells promotes pathogenicity of *K.pneumoniae* by induction of proinflammatory cytokines (21). Proinflammatory cytokines have a protective effect against *K.pneumoniae* by recruitment of neutrophils to the infection site. However, studies pointed out the evasion strategy of virulent *K.pneumoniae* through yersiniabactin secretion(15, 21, 25–27). One important effect of yersiniabactin is evasion from innate immune protein Lipocalin 2 which is produced by neutrophils or mucosal surfaces (25). The other effect of yersiniabactin is the enhancement of bacterial survival in phagocytic cells by reduction of the oxidative stress response (26). We proposed that, despite their high internalization rate by neutrophils, high survival index of the *kfu*^+^-*ybtS*^+^ producing ColR-HvKp could be explained by their resistance inside neutrophils after being uptaken. Capsular polysaccharides of HvKp ST258 were reported to have an inhibition on phagocytosis activity of neutrophils (15). In this study, we did not find a difference in capsule types of ColR-HvKp and other clones.

Another significant finding of our study was NET release from neutrophils after encountering *kfu*^+^*ybtS*^+^ ColR-HvKp (figure 3). We observed the extensive spread of myeloperoxidase and histone in the extracellular space of neutrophils. The role of NETs in the pathogenesis of infection is still under debate. While some pathogens are killed by NETs, others may survive or even benefit from NETs (28). Phagocytosis is a critical event of decisions to form NETs, and if the bacterium is killed by phagocytosis, only a few azurophilic granules may leak into extracellular space with no NET formation (29). Similarly, the *kfu*^-^-*ybtS*^-^ isolates of our study induced very rare NETs with a low amount of myeloperoxidase and histones in extracellular space (figure 3). As the control of our experiment, we studied Klebsiellla ATCC strain, and we did not observe NETosis. Branzk et al. reported that NETs are formed in response to large pathogens. Virulent bacteria may circumvent phagocytosis by the formation of large aggregates and trigger NETosis (29). The successful survival of *kfu*^+^-*ybtS*^+^ isolates (median survival index 7.9) from phagocytosis with induction of extensive NETosis suggested us that protective function of iron uptake systems from being killed by neutrophils might be one of the reasons for mortality of the patients through increased inflammation.

In this study, the 30-day fatality of ST101 and ST395 were 58% and 75%. *K.pneumoniae* ST101 is known as hypervirulent clone mostly responsible for pneumonia and bacteremia in intensive care units. (30, 31). In our study, ColR-HvKP ST101 isolates were found to be associated with VAP infections (p=0.009). In one of our previous studies, 30 day fatality of infections with ST101 *K.pneumoniae* was found to be 72% (30). The ST395 clone is known as a potentially high-risk clone (32), and recent studies pointed out the emergence of carbapenem-resistant ST395 in France and Italy (33, 34). KPC-2 producing ST101 K. pneumoniae was shown to have the highest number of virulence genes associated with capsule type, attachment and iron uptake than the other epidemic clones of *K.pneumonia* (10). Similarly, we observed more virulence genes for iron uptake system, attachment and mucoid phenotype among isolates belonged to ST101 and ST395 than the other clones (heatmap, supplement).

Our novel findings in depiction of pathogenesis of HvKp strains should be supported by the animal studies. Particularly, an animal lung infection model should be developed.

In conclusion, iron uptake systems have a significant contribution to the pathogenesis hypervirulent *K.pneumoniae* ST101 and ST395 infections. These systems enhance successful survival of *K.pneumoinae* against neutrophil phagocytic defense, and stimulate excessive NET formation. The drugs targeted to ferric uptake systems would be a promising approach for treatment of hypervirulent *K.pneumoniae* infections.

## Acknowledgements

Authors declare no conflict of interest. Koc University Institutional Review Board approved the study by the number of 2015.048.IRB1.008. Mehmet Gönen was supported by the Turkish Academy of Sciences (TÜBA-GEBİP; The Young Scientist Award Program) and the Science Academy of Turkey (BAGEP; The Young Scientist Award Program). The authors gratefully acknowledge use of the services and facilities of the Koç University Research Center for Translational Medicine (KUTTAM), funded by the Presidency of Turkey, Presidency of Strategy and Budget. The content is solely the responsibility of the authors and does not necessarily represent the official views of the Presidency of Strategy and Budget.

